# Forces on nascent polypeptides during membrane insertion and translocation via the Sec translocon

**DOI:** 10.1101/310698

**Authors:** Michiel J.M. Niesen, Annika Müller-Lucks, Rickard Hedman, Gunnar von Heijne, Thomas F. Miller

## Abstract

During ribosomal translation, nascent polypeptide chains (NCs) undergo a variety of physical processes that determine their fate in the cell. Translation arrest peptide (AP) experiments are used to measure the external pulling forces that are exerted on the NC at different lengths during translation. To elucidate the molecular origins of these forces, a recently developed coarsegrained molecular dynamics (CGMD) is used to directly simulate the observed pulling-force profiles, thereby disentangling contributions from NC-translocon and NC-ribosome interactions, membrane partitioning, and electrostatic coupling to the membrane potential. This combination of experiment and theory reveals mechanistic features of Sec-facilitated membrane integration and protein translocation, including the interplay between transient interactions and conformational changes that occur during ribosomal translation to govern protein biogenesis.

## Introduction

Co-translational protein biogenesis is tightly regulated to ensure that newly synthesized proteins are correctly targeted and folded within the cellular environment. Throughout this process, a nascent polypeptide chain (NC) is exposed to a complex range of forces and interactions, the study of which is complicated by the crowded, stochastic nature of the cell. The current work combines arrest peptide (AP) experiments and simulation to connect the pulling forces experienced by a NC to the underlying molecular processes associated with membrane integration and translocation via the Sec translocon.

Most membrane proteins and many secretory proteins are targeted to the Sec translocon during ribosomal translation (reviewed in Refs. 1–6). The translocon is a protein-conducting transmembrane channel that is ubiquitous across all kingdoms of life. The ribosome docks onto the cytosolic opening of the translocon, cotranslationally inserting the NC into the translocon channel. The central pore of the translocon facilitates the translocation of hydrophilic loops across the cell membrane, and a lateral gate enables passage of transmembrane domains into the cell membrane [7]. The components of the translocon have been characterized structurally [7–11] and biochemically [12–16], and extensive work has focused on the role of the translocon on regulating NC translocation versus membrane integration [17–23]. Nonetheless, open questions remain about the nature of the transient interactions between the NC and the translocon channel interior and membrane environment.

AP experiments probe the co-translational forces that act on the NC, providing a signature of the underlying interactions between the NC and the translocon during cotranslational membrane integration. Once an AP is synthesized by the ribosome, it stalls further NC translation [24]; the stall is released with a rate that is dependent on the pulling forces that are experienced by the NC [25]. APs are used in nature to control NC translation [24] and have recently been applied to gain insight into physical processes such as integration into the cell membrane [26], co-translational folding [27, 28], and electrostatic interactions [29]. In this study, we use AP experiments with engineered NCs to measure the forces exerted during membrane integration and translocation. To complement the AP experiments, simulations are performed using a recently developed structurally detailed coarse-grained molecular dynamics (CGMD) model that provides nm-lengthscale resolution [30], allowing for the direct computation of the NC dynamics, interactions, and resulting pulling forces. The combination of simulation and experiment elucidates the diverse interactions and forces acting on the NC at specific lengths during translation.

## Results

We consider a series of NC substrates to validate the combined simulation and experimental approach and to investigate the molecular interactions that govern co-translational NC integration and translocation. All NC substrates described in this work utilize a well-established model system, with an engineered domain (H segment) incorporated into the leader peptidase (LepB) protein (Figure 1a, bottom) [17, 26, 29]. We study the forces exerted on the NC during (*i*) the integration of a model transmembrane domain, (*ii*) translocation and integration of non-spanning hydrophobic domains, and (*iii*) the translocation of model hydrophilic and charged domains. CGMD simulations are compared with both previously published [26, 29] and new AP experimental data, providing validation for the computational method and yielding insight into the interactions that govern co-translational NC integration and translocation via the Sec translocon.

**Figure 1.**
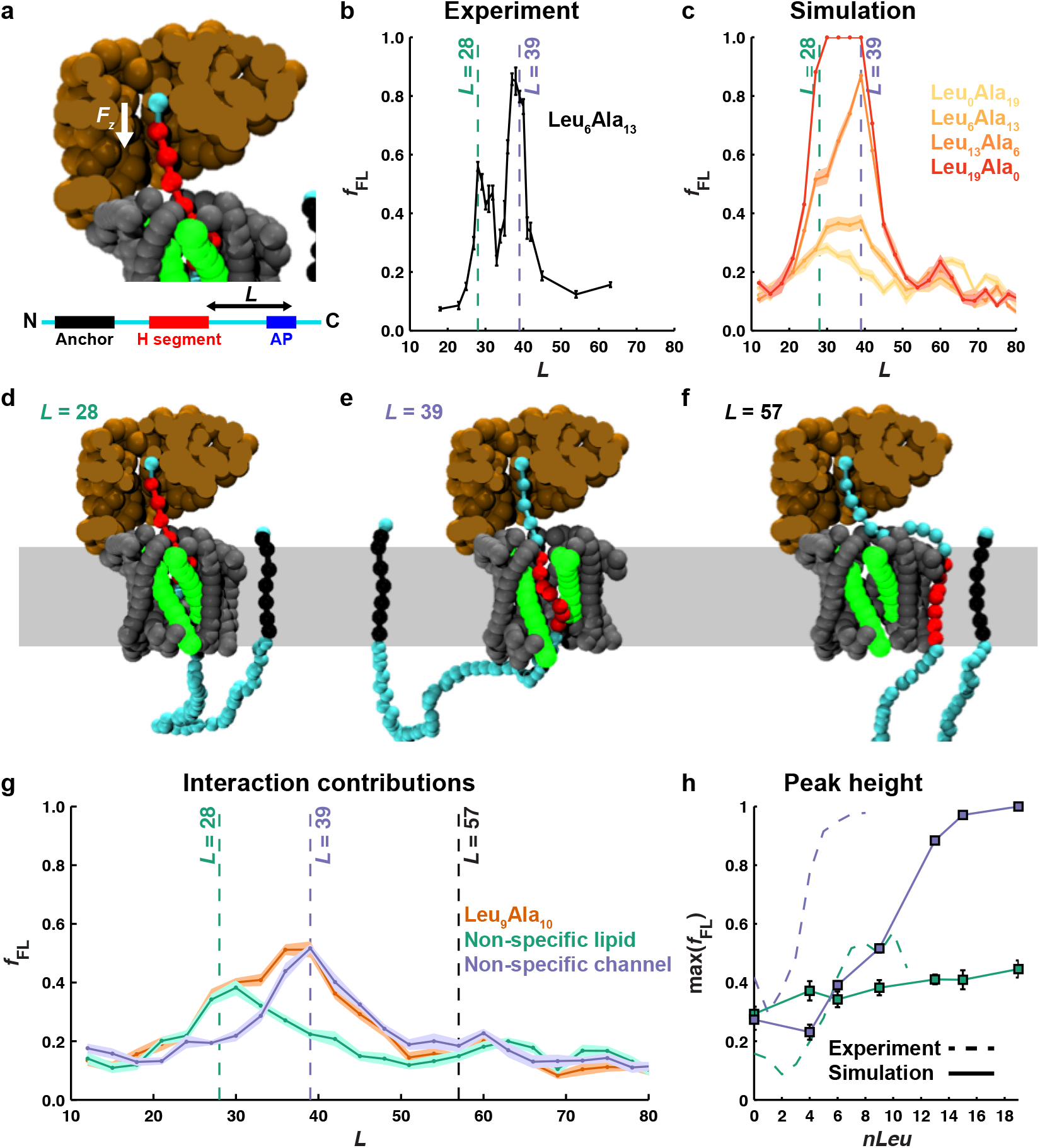
Characterization of the physical processes that drive integration of a hydrophobic transmembrane domain. (**a**) CGMD simulation setup used to calculate pulling forces acting on an engineered hydrophobic H segment (red) during co-translational integration. Shown is a CGMD snapshot at *L* = 28; the C-terminal bead is held fixed and forces exerted by the nascent protein on that bead are calculated. (**b**) Experimental data reproduced from ref [26]. Two peaks in the pulling-force profile are observed during the co-translational integration of the hydrophobic H segment. (**c**) CGMD data for H segments of varying Leucine content. Vertical dashed lines indicate the position of the corresponding peaks in the experimental results. (**d-f**) Representative CGMD configurations at *L* = 28 (d), *L* = 39 (e), and *L* = 57 (f). (**g**) CGMD pulling-force profiles for an H segment with nine leucine residues with default interactions (red), non-specific lipid interactions (green), and non-specific channel interactions (blue). (**h**) The maximum value of *f*_FL_ for the peak near *L* = 28 (green) and *L* = 39 obtained from CGMD (solid lines, as explained in text) and experiment [26] (dashed lines). Error bars indicate the standard error of the mean.

### Forces on integrating hydrophobic domains, and the mechanism of the biphasic pulling force

We begin by investigating the forces of co-translational integration, with the H segment comprised of a model transmembrane domain (Figure 1a, bottom). Previously published AP experiments [26] reveal the points during translation at which increased pulling forces are exerted on the NC (Figure 1b). In these experiments, an AP is inserted downstream of the H segment, and the number of residues between the C-terminal end of the AP and the C-terminal end of the H segment, *L*, is varied (Figure 1a, bottom). The H segment has a fixed length of 19 residues that are either leucine or alanine, and various H segment compositions are tested. The degree of stall-release is experimentally quantified using the fraction of full-length protein, *f*_FL_, as a proxy for the pulling force acting on the AP, with greater forces leading to increased stall-release (see Methods) [25, 26]. Two peaks in *f*_FL_ are observed at *L* = 28 and *L* = 39 (Figure 1b). For comparison with the experiment, CGMD simulations are used to calculate the co-translational forces exerted on the NC for the same sequences as those experimentally tested. The protein sequences are mapped into a coarse-grained representation (Figure 1a, top) and forces acting on the end of the NC that is tethered to the ribosome are directly calculated (Figure 1a, white arrow). Calculated forces are converted to *f*_FL_ assuming Bell’s model to relate force to the force-dependent rate of stall release (see Methods) [25]. The CGMD successfully captures peaks in *f*_FL_ at the same values of *L* (Figure 1c, dashed vertical lines) as previously observed experimentally. Consistent with the experiment, the peaks in *f*_FL_ are dependent on the number of leucine-residues, *nLeu*, in the H segment (Figure 1c).

To identify the physical processes that underlie the observed peaks in both the experimental and simulated pulling-force profiles, we analyze the CGMD trajectories. A characteristic MD configuration for *L* = 28 is shown in Figure 1d. At this NC length, the N-terminus of the H segment first reaches the interior of the translocon, allowing for attractive, residue-specific interactions; this interpretation of the first peak in the pulling-force profile is consistent with experimental data on the effects of point mutations in the H segment [26]. In Figure 1e, a characteristic MD configuration for *L* = 39 is shown. At this point, the N-terminus of the H segment is first able to partition from the interior of the translocon channel into the interior of the lipid membrane via the open lateral gate; this interpretation of the second peak is again consistent with available experimental mutagenesis data [26]. Finally, in Figure 1f, a characteristic configuration associated with larger values of *L* is presented; at these NC lengths, the H segment has completed integration into the lipid membrane and the NC is no longer under tension.

The molecular origin of the observed peaks is further confirmed by additional CGMD simulations with modified interactions. Considering first the H segment with nine leucine residues, Figure 1g shows the pulling-force profile calculated from simulations for which (blue) the residue-specificity of the interactions between the NC and the translocon are eliminated or (green) the residue-specificificity of the water-lipid transfer free energies are eliminated (see Methods). The simulations without residue-specific interactions between the NC and the translocon channel do not display the pulling-force peak at *L* = 28, confirming that the first peak reports on the specific interactions between the H-domain and the residues of the translocon interior. Similarly, the simulations without residue-specific water-lipid transfer free energies do not display the pulling-force peak at *L* = 39, confirming that this second peak arises from the partitioning of the H-domain from the translocon interior into the membrane interior. Similar results are obtained for all tested H segments, with the pulling-force profiles consistently comprised of two underlying peaks (Figure S1). These results provide clear validation of the CGMD simulations in comparison to experiment, as well as direct evidence of the physical origins of the observed features in the pulling-force profiles.

Finally, to examine the dependence of the pulling forces on the sequence of the H segment, we compare the peak heights in the pulling-force profiles, max(*f*_FL_), for H segments with varying leucine content. The peak heights are calculated separately for the peak near *L* = 28 and the peak near *L* = 39, using simulations with modified interactions (as in Figure 1g). Figure 1h presents the peak heights as a function of the number of leucine residues in the H segment, calculated from trajectories without specific interactions between the NC and the translocon (blue) or without specific interactions between the NC and the lipid membrane (green). The figure shows that with increasing number of leucine residues, *nLeu;* the peak associated with the NC-translocon interactions (green) remains unchanged, while the peak associated with NC-lipid interactions increases. Experimentally determined peak heights are shown as dashed lines for comparison (*L* = 28 in green and *L* = 39 in blue). Although CGMD underestimates the height of peaks in the pulling-force profile as compared to the experiment, the hydrophobicity dependence and the reduced sensitivity of the peak at *L* = 28 are reproduced.

### Forces on hydrophobic domains of variable length

To examine the relation between the size of the hydrophobic domain and the forces exerted on NC, we investigate hydrophobic poly-leucine H segments of varying length using both CGMD simulations and new AP experiments. This “variable-length assay” allows for comparison of short hydrophobic domains (illustrated in Figure 2a) that primarily undergo membrane translocation versus longer hydrophobic domains that primarily undergo membrane integration; it contrasts with the “fixed-length assay” from the previous section in which all H segments were the same length and sufficently long to span the membrane.

**Figure 2.**
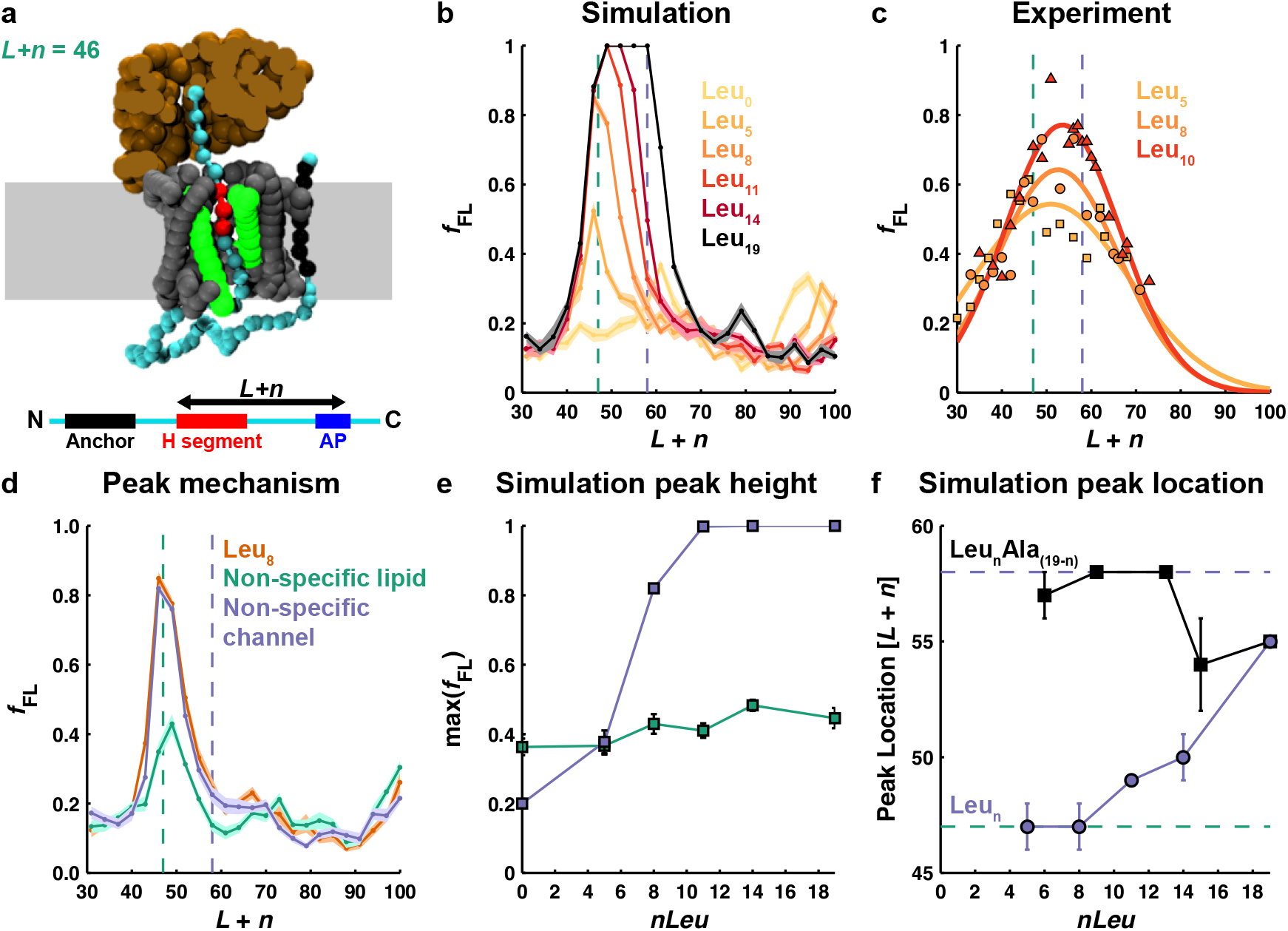
Forces exerted on hydrophobic domains of variable length. (**a**) CGMD snapshot for an H segment with eight leucine residues (red), stalled at *L* + *n* = 46. The pulling-force profile determined from CGMD (**b**) and from experiment (**c**) for poly-leucine H segments with increasing numbers of leucine residues. Scatter points reflect experimental data-points, the solid lines are a single-gaussian fit of the experimental data-points. (**d**) CGMD pulling-force profile for an H segment with eight leucine residues with default interactions (red), non-specific lipid interactions (green), and non-specific channel interactions (blue). (**e**) The maximum value of *f*_FL_ from CGMD in which the peaks were isolated as shown (d). (**f**) Location of the lipid-interaction peak in the CGMD pulling-force profile as a function of *nLeu*. For poly-leucine H segments (blue) and for 19-residue H segments consisting of alanine and leucine (black). The dashed lines correspond to the *L* + *n* value at which the channel interaction peak (green) and the lipid interaction peak (blue) are observed for the fully spanning transmembrane domains in the fixed-length assay. Error bars indicate the standard error of the mean.

Figures 2b and c present pulling-force profiles for the poly-leucine H segments of varying length, n, obtained using CGMD and experiments, respectively. Results are plotted as a function of the length of the NC chain from the C-terminus of the AP to the N-terminus of the variable-length H segment, *L* + *n*, where *L* is defined as before (Figure 2a, bottom). With this choice for the x-axis, the expected position for the peaks associated with the NC-translocon interactions and the NC-lipid interactions from the fixed-length assay in the previous section (green and blue vertical lines, respectively) are independent of the variable length of the H segment. Both simulation and experiment show a single broad peak in the pulling-force profile (Figures 2b and c), compared to the two distinct peaks observed for model transmembrane domains in Figure 1. With increasing length of the hydrophobic domain, the observed single peak broadens and increases in height.

To deconvolute the role of NC-translocon versus NC-lipid interactions in Figures 2b and c, CGMD simulations with modified interactions are performed, as before. Considering first the H segment with eight leucine residues, Figure 2d contrasts the results obtained using non-specific lipid interactions (green) versus non-specific translocon interactions (blue). Consistent with the fixed-length assay (Figure 1g), the simulations with non-specific lipid interactions (green) yield a peak at the expected NC length due to residue-specific interactions between the NC and the translocon. However, the simulations in Figure 2d with non-specific translocon interactions (blue) yield a peak at shorter NC lengths than expected from the fixed-length assay. Figure S2 presents the analog of Figure 2d for the NC sequences with different H-domain length. To further validate the deconvolution of the pulling-force profile in Figure 2d into two distinct peaks with different physical origins, Figure 2e presents the calculated peak heights across the various H-domain lengths as a function of the number of leucine residues, *nLeu*, revealing a trend that is consistent with the fixed-length assay (Figure 1h); specifically, the peak height associated with the NC-translocon interactions (green) remains unchanged, while the peak height associated with NC-lipid interactions increases. Finally, Figure 2f contrasts the position of the peak associated with NC-lipid interactions from the variable-length assay (blue) versus the corresponding results from the fixed-length assay from the previous section (black). In contrast to the fixed-length assay, for which the NC-lipid peak position is relatively invariant with respect to the increasing number of leucine residues, the results from the variable-length assay find that the peak position steadily increases with the number of leucine residues.

The contrasting behavior of the fixed-versus variable-length assays in Figure 2f provides insight into the mechanism by which hydrophobic portions of the NC sample the lipid membrane, an issue that has been the focus of considerable discussion [4, 31–35]. Observation of the peak in the pulling-force profile associated with the NC-lipid interaction requires that the lateral gate of the translocon be in the open conformation, to allow for contact of the NC with the lipid environment. Previous work has suggested that opening of the translocon lateral gate is stabilized when hydrophobic NC residues reside in the translocon channel interior [34, 35]. Note that for a given number of leucine residues, *nLeu*, the variable-length assay prescribes that those hydrophobic residues appear consecutively in the H-domain sequence, whereas the fixed-length assay dilutes the hydrophobic leucine residues over a total of 19 residues. If a threshold number of hydrophobic residues is needed to stabilize the free-energy of opening of the translocon lateral gate [35], then the variable-length assay will reach that threshold at shorter lengths of the NC than the fixed-length assay. Consistent with this mechanism for lateral gating, the lateral gate in the CGMD is found to open at shorter NC lengths for the H segments used in the variable-length assay (Figure S3). This also explains why the lipid-interaction peak for small values of *nLeu* appears at shorter NC lengths in the variable-length assay than in the fixed-length assay, as well as why the lipid-interaction peak position increases for the variable-length assay but not for the fixed-length assay. The experimental and simulation results presented in Figure 2 thus provide evidence in support of the hydrophobic stabilization of the open-state state for the lateral gate of the translocon [35], as well as the prediction that the H-domain samples the membrane environment across the lateral gate as it passes down the axis of the translocon channel [36, 37]; it is likewise consistent with the “sliding” model for transmembrane helix integration, which posits that hydrophobic segments in the NC slide along the lateral gate of the translocon, with one side exposed to lipid [4].

### Pulling forces on hydrophilic domains

Previously published AP experiments indicate that significant pulling forces act on hydrophilic segments of the NC during translocation in *E. coli* [29]. These forces were attributed to the coupling of negatively charged residues on the NC with the membrane electrostatic potential. Here, we investigate this underlying mechanism using CGMD, finding broad agreement with the previously proposed mechanism, as well as identifying additional features in the pulling-force profiles that are attributed to interactions of charged residues in the NC with the charges on the ribosome and to changes in the NC solvation environment.

Figure 3a presents pulling-force profiles for three distinct hydrophilic H segments (D_5_, Q_5_, and K_5_) obtained from previous AP experiments [29]. Results are plotted as a function of *L* + *n*, with *n* = 5 for all considered cases. A dominant peak at *L* + *n* ≈ 50 in the pulling-force profile is observed for the negatively charged D_5_ H segment (red); the peak was found to reduce in magnitude, in a concentration dependent manner, when indole was added to the growth medium, which suggests that the peak is due to the membrane electrostatic potential [29]. The corresponding peak in the pulling-force profile is not found for the charge-neutral H segment (Q_5_, green) nor the positively charged H segment (K_5_, blue). Interestingly, the negatively charged H segment also exhibits a somewhat larger value for the pulling force at shorter NC lengths (*L* + *n* < 45) in comparison to the other sequences; this feature was found to recur in a variety of negatively charged NC sequences (Figure S4a of Ref. 29) although a theoretical explanation for the feature was not provided.

**Figure 3.**
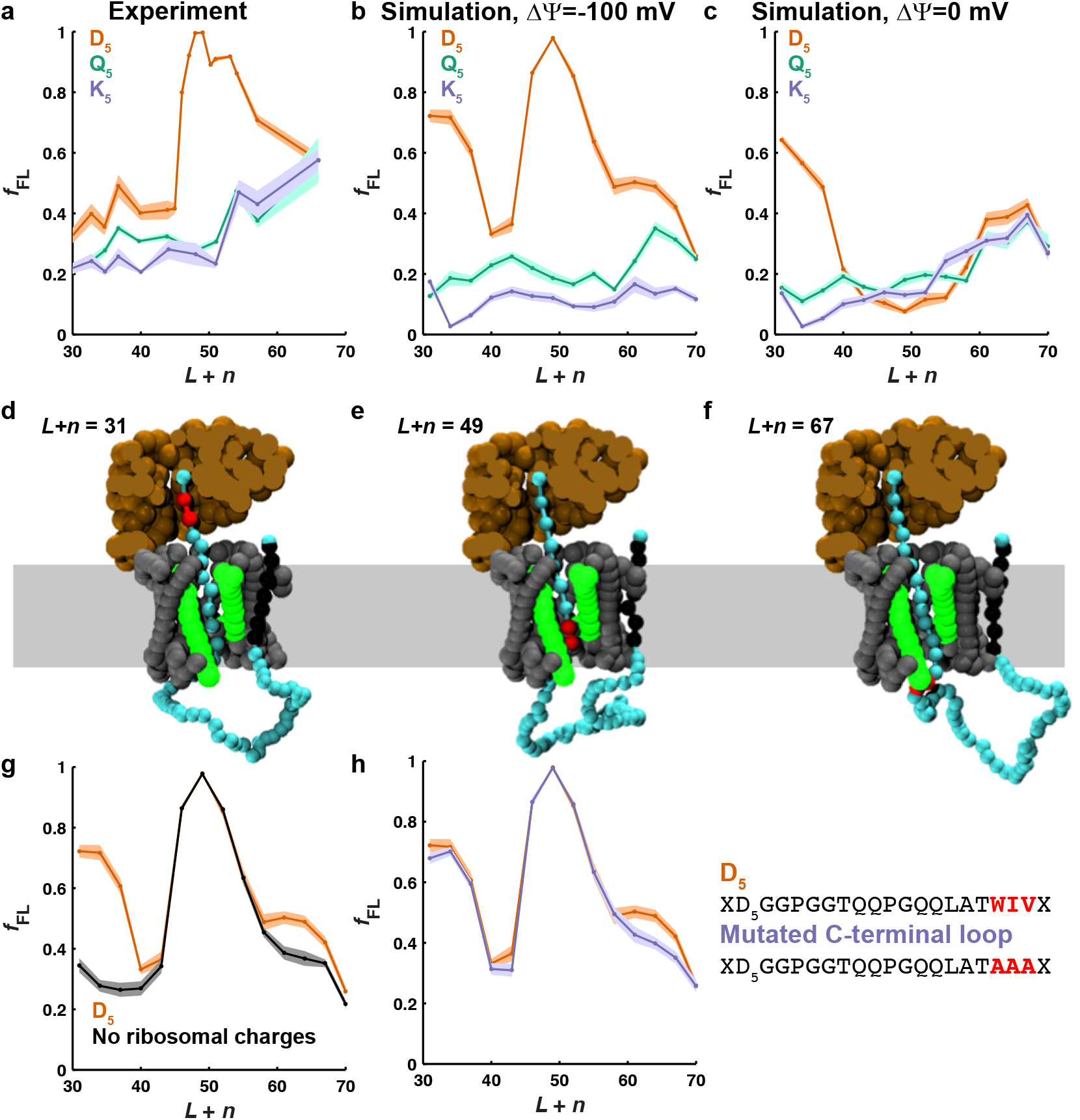
Forces exerted on hydrophilic H segments. The pulling-force profile determined from experiment [29] (**a**) and from CGMD (**b**) for negatively charged (D_5_, red), positively charged (K_5_, blue), and neutral (Q_5_, green) 5-residue H segments. (**c**) As in (**b**), but for CGMD simulations without a membrane potential. (**d-f**) CGMD snapshot for a D_5_ H segment (red), stalled at; *L* + *n* = 31 (**d**), *L* + *n* = 49 (**e**), and *L* + *n* = 67 (**f**). (**g**) The pulling-force profile for a D_5_ H segment with (red) and without (black) ribosomal charges. (**h**) The pulling-force profile for a D_5_ H segment with the original C-terminal loop (red) and with a mutated C-terminal loop that is more hydrophilic (blue). Error bars indicate the standard error of the mean.

To explore the mechanistic origin of these pulling-force features, CGMD pulling-force profiles are obtained using the same protein sequences, calculated with (Figure 3b) and without (Figure 3c) the approximate E. coli membrane potential (ΔΨ = −100 mV). For the CGMD results obtained in the presence of the membrane potential, the calculated pulling-force profiles are in good agreement with experiment, showing a dominant peak at *L* + *n* ≈ 50 for the negatively charged H segment and no such features for the other H segments; additionally, it is seen that the pulling-force profile for the negatively charged H segment at short NC lengths (*L* + *n* < 45) is increased in comparison to the other sequences, albeit to a greater degree than is observed experimentally. Figure 3c shows that removing the membrane potential in the CGMD simulations leaves all features of the pulling force unchanged, except for the dominant peak in the D_5_ profile (red) at *L* + *n* ≈ 50. The membrane-potential sensitivity of the D_5_ peak at *L* + *n* ≈ 50 is in good agreement with the previous experimental studies of indole concentration dependence [29]. From both CGMD and experiment, these results suggest that the dominant D_5_ peak at *L* + *n* ≈ 50 arises from coupling of the negatively charged residues to the membrane potential, whereas a different mechanism leads to the greater pulling forces on D5 for shorter NC lengths in comparison to the other H segments.

To illustrate the interactions of the H segment at various NC lengths, Figures 3d-f present snapshots of the CGMD simulations for the sequence with the D_5_ H segment (indicated in red beads). At short NC lengths (part d), the H segment remains in close proximity to the ribosome, and it does not extend to the membrane interior regions where the membrane potential significantly varies; this is consistent with the finding that the membrane potential exhibits minimal forces on the H segment at these NC lengths. For NC lengths associated with the dominant peak in the D_5_ pulling force profile at *L* + *n* ≈ 50 (part e), the H segment extends to the membrane interior, where the membrane potential is most rapidly varying and will exert the largest pulling forces on the negatively charged residues. Finally, for even larger NC lengths (part f), the hydrophilic H segment is fully translocated across channel and is favorably solvated in the hydrophilic environment of the periplasm.

While the CGMD pulling-force profiles (Figures 3b and c) and the simulation snapshots at *L* + *n* ≈ 50 (Figure 3e) are completely consistent with the interpretation that the dominant peak in the D5 pulling force profile is due to the membrane potential, the CGMD simulations provide additional insight into the mechanistic features of the pulling-force profile at both shorter and longer NC lengths. In particular, at shorter NC lengths (*L* + *n* < 45), it is observed in both CGMD and experiment that D_5_ exhibits larger pulling forces than the other two H segments. The simulation snapshot at Figure 3d suggests that this feature in the D5 profile may arise from repulsive electrostatic interactions between the negatively charged NC and the negatively charged ribosomal RNA. Figure 3g tests this hypothesis using the CGMD model, comparing the original D_5_ pulling-force profile (red) with that obtained in the absence of charges on the ribosome (black); clearly, the pulling force profile at short NC lengths in the absence of ribosomal charges returns to the baseline of the other H segments, supporting the hypothesis.

Finally, we investigate the rise in the pulling force profile that is observed for all three hydrophilic H segments at long NC lengths (*L* + *n* > 60), in both experiment (Figure 3a) and in the CGMD simulations (Figure 3b and even more clearly in Figure 3c). This feature is relatively independent of the charge of the hydrophilic domain; it was therefore not explained by previous work that focused on the role of charged residues [29]. From the CGMD snapshot in Figure 3f, it is clear that the H segment has extended beyond the translocon interior at these NC lengths, such that the H segment has been replaced by C-terminal residues of the NC in the channel interior. This suggests a hypothesis in which the observed pulling forces at long NC lengths is due to the favorable free energy associated with transferring the hydrophilic H segment from the amphiphilic (or weakly hydrophilic) interior of the channel to the strongly hydrophilic environment of the periplasm. To test this hypothesis, Figure 3h compares the original D5 pulling-force profile (red) with that obtained by increasing the hydrophilicity of the residues in the C-terminal tail (C-tail) of the NC, thus counterbalancing the favorable free energy of transferring the D5 from the channel interior to the periplasm with an unfavorable free energy of transferring the C-tail from the hydrophilic cytosol the channel interior. This alteration of the C-tail sequence leads to a reduction of the pulling force profile at long NC lengths, confirming that the the increased pulling forces at long NC lengths arises from free energy of transferring the hydrophilic H segment from the channel interior to the periplasm.

## Discussion

Membrane integration and protein translocation via the Sec translocon are critical steps in the biosynthesis and targeting of proteins in cells. The current work probes the fundamental interactions and conformational changes associated with these processes, using AP experiments to measure the pulling forces associated with interactions of the NC with its environment [25, 38]. We present new AP experiments to obtain the pulling-force profiles as a function of NC length during translation, as well as detailed analysis of these and previously reported AP experiments [26, 29] using long-timescale CGMD simulations [30] that allow for the direct computation of the pulling-force profiles.

Engineered NC sequences allow for the investiation of co-translational forces that act on transmembrane hydrophobic domains (Figure 1), non-membrane-spanning hydrophobic domains (Figure 2), and translocating hydrophilic domains (Figure 3). For each NC sequence, experimental pulling-force profiles are directly compared with those obtained using CGMD, validating the simulation method. Analysis of the microscopically detailed CGMD simulations provides insight into the mechanistic origins for the experimentally observed features pulling-force profiles.

Several conclusions emerge from this work. Firstly, a detailed analysis of the mechanistic origin of biphasic pulling-force profiles for transmembrane hydrophobic domains is provided [26]; deconvolution of the pulling-force profiles using the CGMD (Figure 1) confirm that the peak at shorter NC lengths arises from translocon-NC interactions, while the peak at longer lengths is associated with NC-lipid interactions during membrane integration. Secondly, consideration of hydrophobic domains of variable length (Figure 2) elucidates the effect of hydrophobic domains on the conformational state of the translocon. The combined experimental and theoretical analysis confirms predictions that hydrophobic segments of the NC stabilize the open state of the tranlsocon lateral gate [35], as well as that even during translocation, sufficiently hydrophobic segments sample the membrane interior as they pass down the axis of translocon channel [4, 37]. Finally, investigation of translocating hydrophilic domains (Figure 3) using CGMD confirms that the dominant peak in the pulling-force profile arises from the coupling of charged residues to the membrane electrostatic potential [29]; however, the CGMD additionally suggest that previously unexplained features of the pulling-force profiles arise from electrostatic repulsion between negatively charged residues in the NC and the ribosomal RNA (at short NC lengths) and from forces associated with partitioning of hydrophilic segments of the NC from the translocon channel interior to the more hydrophilic environment of the periplasm (at long NC lengths).

Taken together, the results presented here demonstrate that AP experiments – combined with long-timescale CGMD simulations that enable the interpretation and deconvolution of the experimentally observed pulling-force profiles – provide rich detail on the interactions and conformational changes associated with Sec-facilitated membrane integration and protein translocation.

## Methods

### Experimental methods

All plasmids were designed as in [26] i.e., H segments of different amino acid composition and flanked by “insulating” GPGG….GGPG segments were inserted into the periplasmic P2 domain of the E. coli inner membrane protein LepB. The 8-residue sequence HAPIRGSP from the Mannheimia succiniciproducens SecM protein [26] was inserted at varying distances downstream of the C-terminal end of the H segment, leaving a 23-reside C-terminal tail after the AP to ensure that arrested and full-length protein products were of sufficiently different molecular weight to allow separation by SDS-PAGE. Constructs with poly-leucine H segment of composition 5L, 8L and 10L were expressed in E. coli, and analyzed by pulse-labeling, immunoprecipitation, and SDS-PAGE as described in ref [26]. The fraction full-length protein, *f*_FL_, was calculated as *f*_FL_ = *I*_FL_/(*I*_FL_+*I_A_*), where *I*_FL_ and *I*_A_ are the intensities of the bands corresponding to, respectively, the full-length and arrested forms of the protein on the SDS-PAGE gel.

### Computational methods

We employ a recently developed CGMD [30] to measure co-translational forces acting on a NC during Sec-facilitated integration into the lipid membrane or translocation across the lipid membrane. CGMD calculates the dynamics of the NC at a μs time-resolution and a nm length-resolution, with explicit ribosomal translation and lateral gating of the Sec translocon [30]. The only modification of the CGMD method from previous work [30] is the inclusion of a membrane electrostatic potential, as described below.

### CGMD Method

For a given NC, the protein sequence is mapped into the coarse-grained (CG) representation (Figure 1a, top), with one CG bead representing three amino-acid residues [30]. NC beads interact with CG beads representing the translocon and ribosome via pair-wise interaction that depend on the charge and hydrophobicity of the NC bead, and the charge and location of the translocon bead. The full geometric coordinates for the CG representation of the ribosome, translocon, and membrane environment are provided in Ref. 30. The interaction between the CG beads and the lipid membrane is accounted for by a water-lipid transfer free energy assigned to each CG bead, derived from the Wimley-White water-octanol transfer free energy [39] of the underlying amino-acid residues. Overdamped Langevin dynamics for the CG beads is simulated with an isotropic diffusion coefficient of 253.0 nm^2^/s and a timestep of 300 ns. The lateral gate of the translocon occupies two discrete conformations (closed and open), with stochastic transitions between the conformations governed by the free-energy difference between the two states as a function of the NC configuration [30].

To simulate AP stalling, translation is halted after a given number of CG beads have been translated. The length *L* of the NC upon stalling corresponds to the number of amino-acid residues counted from the C-terminus of the H segment to the C-terminus of the AP (Figure 1a). The instantaneous pulling force on the NC, *F*_z_, is calculated as the component of the force along the translocon channel axis that acts on the most C-terminal bead in the CGMD; this C-terminal bead is held fixed to mimic translation arrest (Figure 1a, top). The value of *L* in the CGMD is then defined as the number of residues that are explicitly represented as CG beads plus a constant correction of 27 residues, accounting for the amino-acid residues between the most C-terminal bead in the CGMD and the end of the AP.

Simulations without residue-specific interactions (Figures 1g and 2d) are performed in exactly the same manner, except with modified interactions for the CG beads. Instead of having interaction parameters as based on the underlying amino-acid sequence [30], parameters are set to a constant value irrespective of the amino-acid sequence. Specifically, for simulations without specific NC-translocon interactions, all NC beads interact with the translocon channel using the parameters for a QQQ tri-peptide (*λ*_c_ = 0.75 and *λ_o_* = 0.78); and for simulations without residue-specific lipid interactions, all NC beads employ a water-lipid transfer free energy of 5*ε*.

### Inclusion of the membrane potential

To investigate coupling of the charged residues in the NC to the membrane electrostatic potential in *E. coli* (Figure 3), a membrane potential is included in the CG model for the calculations reported in the section *Pulling forces on hydrophilic domains*. Following previous work [29], the potential energy function, *U*_mp_, associated with the additive interaction of the membrane potential with each charged CG bead, *i*, is described using

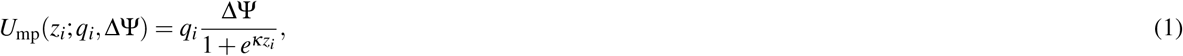

where *z_i_* is the position of the bead along the channel axis, *q_i_* is the charge of the bead, *κ* = 1.6σ^−1^ (0.2Å^−1^) is the reciprocal lengthscale of the membrane potential drop, and ΔΨ is the value of the maximum potential drop of −3.74*ε* (−100 mV) used for the results in Figure 3b, g, and h or 0*ε* (0 mV) used elsewhere.

### Calculation of fraction full-length protein

To compare with the available experimental data, the pulling forces calculated from the CGMD must be converted to a prediction of the fraction of full-length protein, *f*_FL_, observed in experiment. Following previous work [25, 29], the AP-stalled ribosome is assumed to restart translation with a force-dependent rate, *k*_FL_, which is calculated assuming Bell’s model,

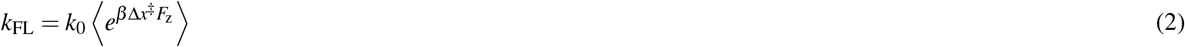

where *k*_0_ is the rate without an applied force, *β* = 1/*k*_B_*T*, Δ*x*^‡^ is an AP-dependent characteristic distance, *F_z_* is the previously defined instantaneous pulling force on the NC, and 〈..〉 indicates ensemble averaging over the CGMD trajectory data. The employed value of Δ*x*^‡^=0.5 nm for all sequences is based on previous work [25, 29]; the value for *k*_0_ is described below.

The ensemble average in Eq. 2 is obtained from CGMD sampling trajectories, with the NC stalled at a given length *L*. Each sampling trajectory is of length 15 s. For each NC sequence and each value of *L*, the ensemble average is obtained by averaging from 100 independent sampling trajectories.

The force dependent rate for breaking the translation arrest, *k*_FL_, is then used to calculate the experimentally observable fraction of full-length protein, *f*_FL_, using

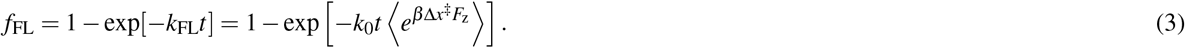

The only undetermined parameter in this equation is (*k*_0_*t*), which depends on the details of the AP, the background pulling-force in the experimental system, and the observation time. We determine a value for (*k*_0_*t*) in this work by fitting the calculated baseline of *f*_FL_ to that observed in experiment (using the data in Figure 1c for *L* >= 51). We emphasize that this fit is done once, yielding a value of(*k*_0_*t*) = 3.7 * 10^−12^ that is held fixed for all other reported results.

The CG mapping of three amino-acid residues to a single CG bead allows for three possible frame-shifts between the amino-acid and CG sequences. At each reported value of *L, f*_FL_ is separately calculated for all three possible frame-shifts (with associated lengths *L* – 1, *L*, and *L* + 1) and the result is averaged. When comparing to experimental data at a given *L*, we refer to the CGMD results for which *L* equals the nearest multiple of three (i.e., experiments at *L* = 28 are compared to CGMD simulations for which the middle frame is *L* = 27). Altogether, 4500 s of CGMD simulation time is performed for each reported value of *f*_FL_.

Note that the relation in Eq. 3 assumes a first-order kinetic scheme in which AP stalling is overcome with a force-dependent rate *k*_FL_, with no off-target pathways. We also considered a slightly more complex kinetic scheme in which the stalled ribosomes experience an additional degradation pathway with a fixed rate; this more complex scheme led to no substantial changes in the results.

## Acknowledgements

GvH acknowledges support from the Knut and Alice Wallenberg Foundation, the Swedish Cancer Foundation, and the Swedish Research Council. TFM acknowledges support from the National Institutes of Health (R01GM125063) and the Office of Naval Research (N00014-16-1-2761). Computational resources were provided by the National Energy Research Scientific Computing Center (NERSC), a DOE Office of Science User Facility supported by the Office of Science of the U.S. Department of Energy under Contract No. DE-AC02-05CH11231 and the Extreme Science and Engineering Discovery Environment (XSEDE) [40], which is supported by National Science Foundation grant number ACI-1548562.

## Supplemental Figures

**Figure S1.**
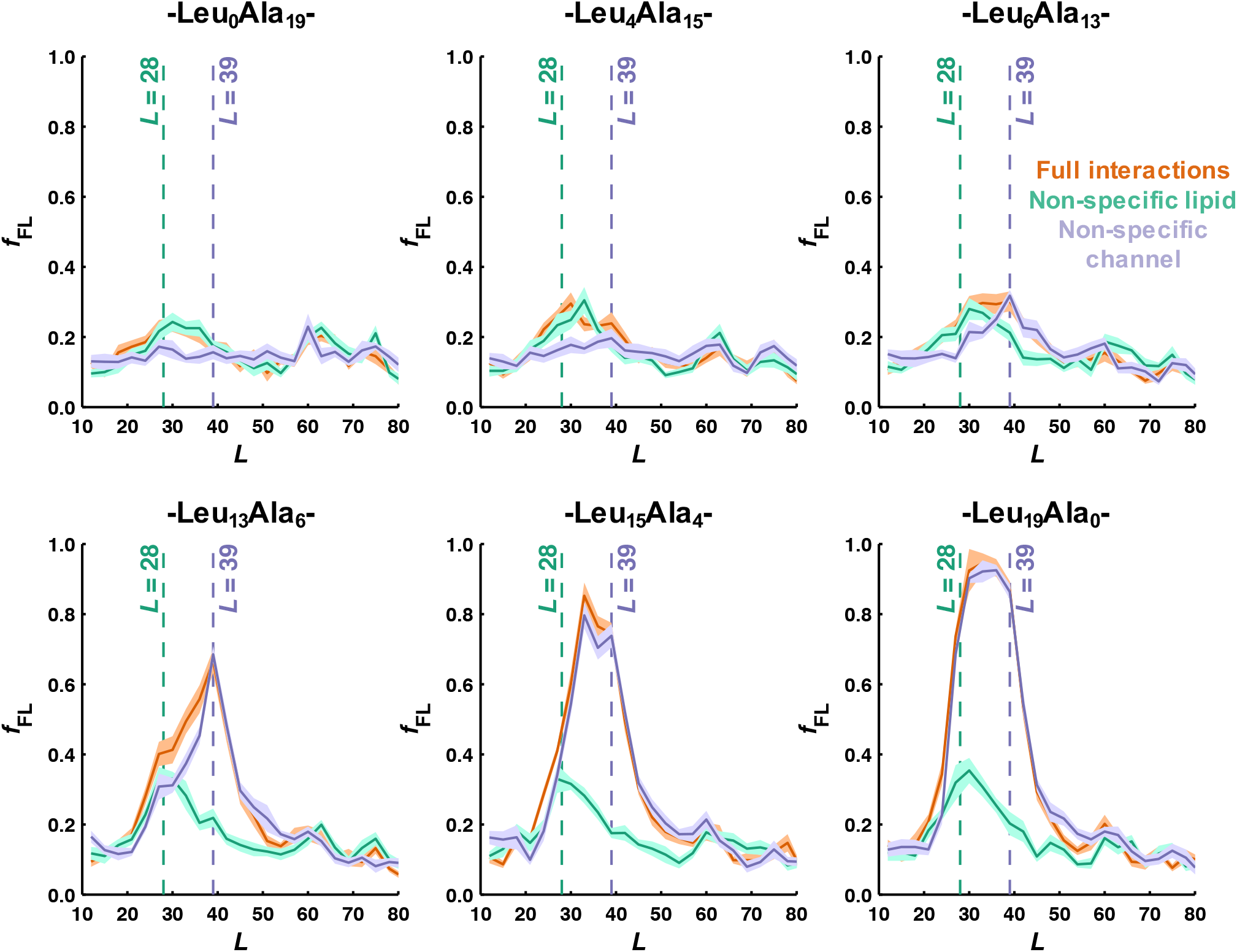
CGMD pulling-force profiles for H segments with varying leucine content (as labeled) with default interactions (red), non-specific lipid interactions (green), and non-specific channel interactions (blue). This data shows that the two separable peaks observed in Figure 1g are observed for all tested H segments. The maximum values in these data were used to construct Figure 1h. Error bars indicate the standard error of the mean.

**Figure S2.**
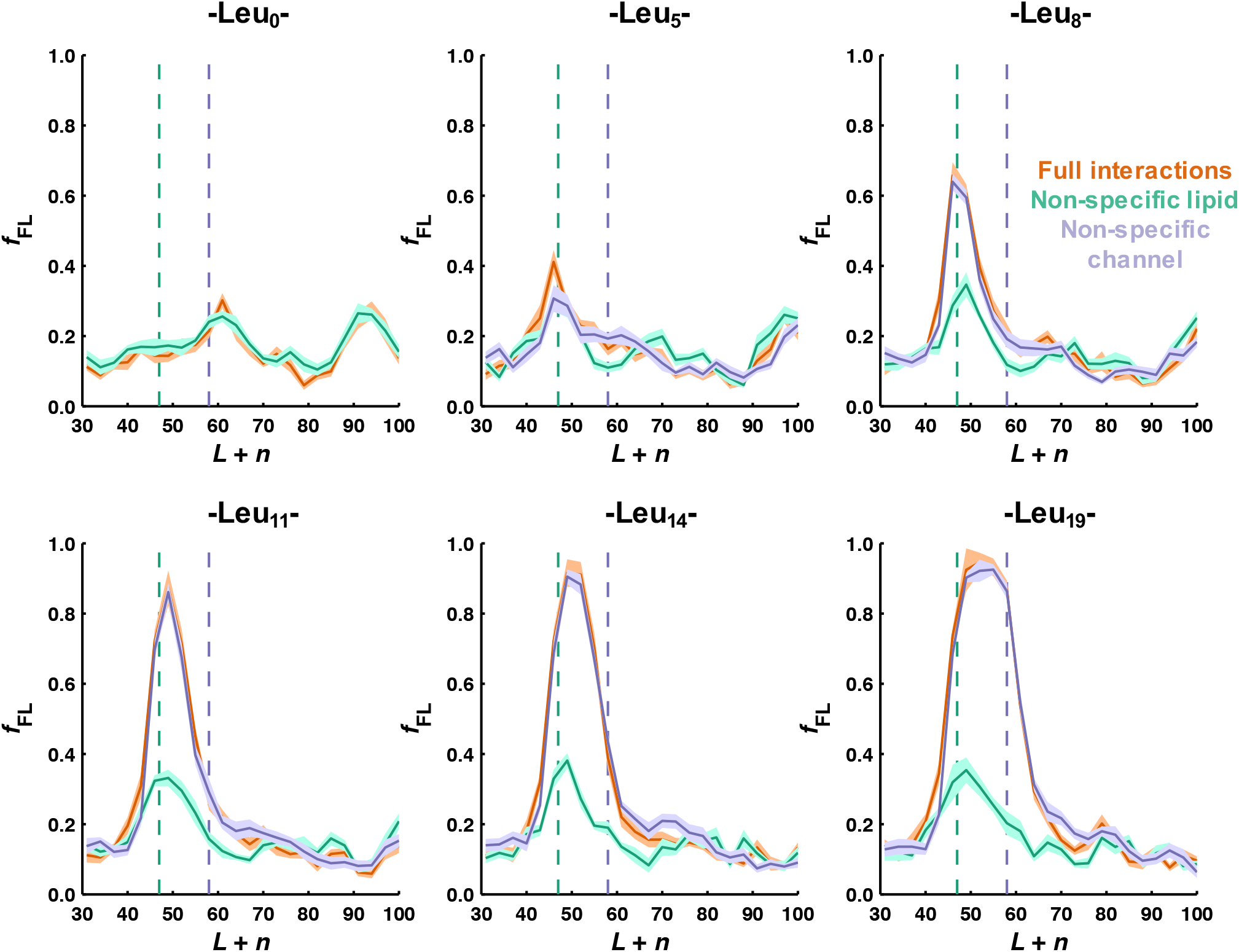
CGMD pulling-force profiles for poly-leucine H segments with varying leucine content (as labeled) with default interactions (red), non-specific lipid interactions (green), and non-specific channel interactions (blue). This data shows that the two separable peaks observed in Figure 2d are observed for all tested H segments. The maximum values in these data were used to construct Figure 2e, and the value of *L* + *n* at which the maximum in *f*_FL_ occurs is reported in Figure 2f. Error bars indicate the standard error of the mean.

**Figure S3.**
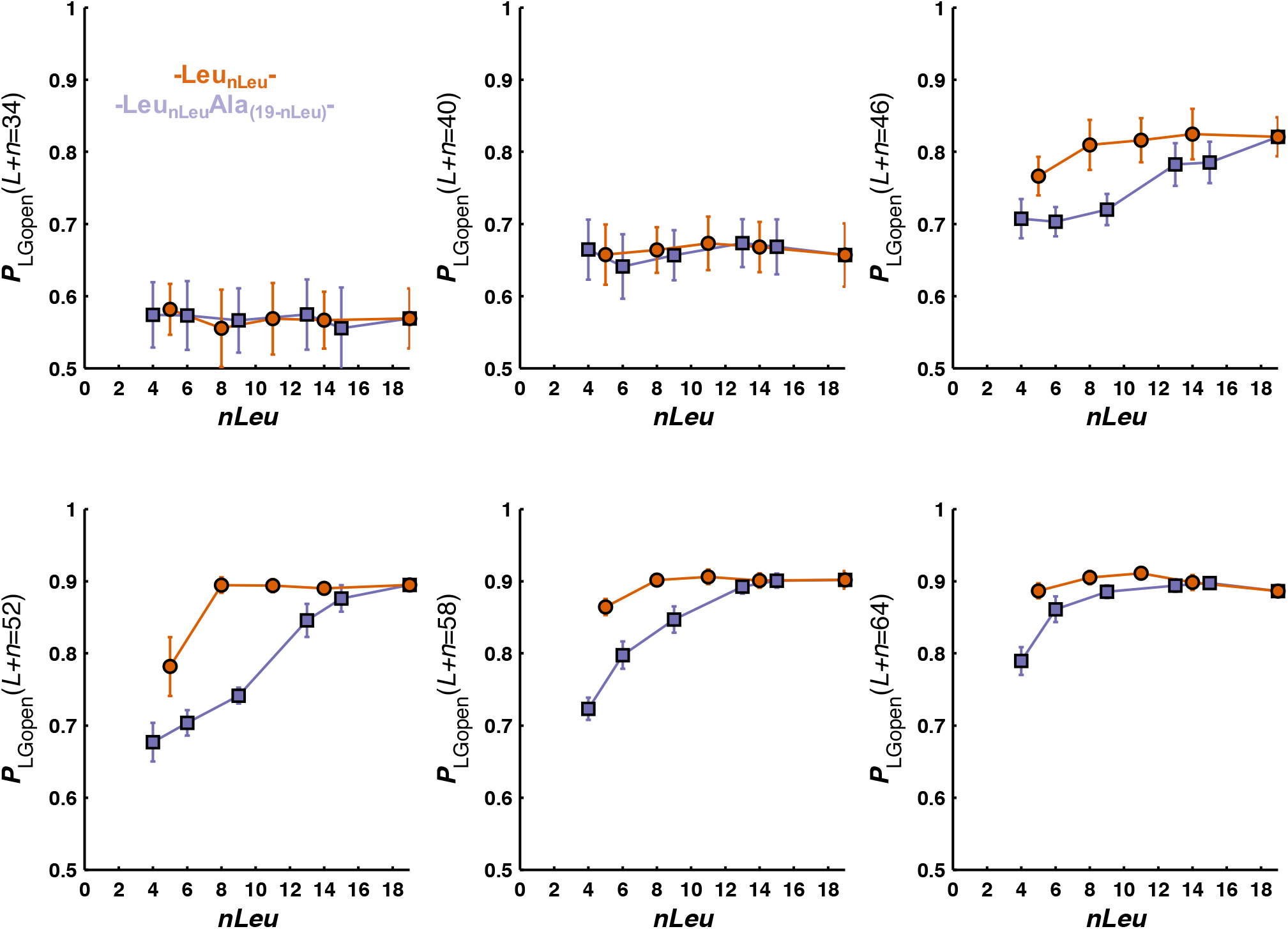
The probability that the lateral gate is in the open conformation, *P*_LGopen_(*L* + *n*), plotted for various values of *L* + n. Data is shown for both poly-leucine H segments (red) and 19-residue H segments consisting of leucine and alanine residues (blue). At values of *L* + *n* corresponding to NC lengths at which the H segment has just reached the translocon (*L* + *n*=46) the lateral gate is more likely in the open conformation for the hydrophobic poly-leucine H segments (red), allowing the H segment to sample the lipid membrane and resulting in a lipid interaction peak that occurs at lower *L* + *n* values (Figure 2f). Error bars indicate the standard error of the mean.

